# The 21-base pair deletion mutant Calpain 3 does not inhibit wild-type Calpain 3 activity

**DOI:** 10.1101/2023.10.03.560718

**Authors:** Swati Maitra, Seungjae Oh, Yun-Jeong Choe, JiHye Kim, Nam Chul Kim

## Abstract

**Introduction:** Calpain 3 is one of the calpain protease family members, which is a calcium-dependent proteolytic enzyme predominantly expressed in skeletal muscle. Loss-of-function mutations in the Calpain 3 gene have been related to autosomal recessive *Limb-Girdle Muscular Dystrophy 1* (LGMDR1), a common form of muscular dystrophy. Recently, the heterozygous 21-bp deletion mutation of the Calpain 3 gene has been reported to cause autosomal dominant *Limb-Girdle Muscular Dystrophy 4* (LGMDD4). According to its dominant inheritance pattern, it has been suggested that the deletion mutant proteins act in a dominant-negative manner. Therefore, we examined whether the mutant protein can suppress the activity of wild-type Calpain 3 and has any dominant toxicity in cell culture and *in vivo Drosophila* models.

**Methods:** A human cell culture (HeLa cells) model with the transient transfection of human wild-type and mutant Calpain 3 and *in vivo Drosophila* models overexpressing wild-type and mutant *Drosophila* Calpain A and B were utilized in this study to assess dominant effects of Calpain 3 21-bp deletion mutant. Western blot analysis was used to determine protein stability and catalytic activity in cell culture. External eye morphology and muscle integrity were examined to observe dominant toxicity in *Drosophila* models.

**Results:** The 21-bp deletion mutation of Calpain 3 resulted in catalytic inactivation, which did not inhibit wild-type Calpain 3 autolytic and catalytic activity against Calpastatin in HeLa cells. In addition, the mutant protein was normally processed by wild-type Calpain 3. Overexpression of wild-type and deletion mutant Calpain 3 in the *Drosophila* eye and muscles did not exhibit significant developmental and age-related dominant toxicity.

**Discussion:** We provide evidence that mutant Calpain 3 does not suppress wild-type Calpain 3 activity. Rather, it is a mutant lacking autocatalytic processing activity like many other loss-of-function Calpain 3 mutants causing LGMDR1. Our results implicate that the stability of the heteromeric mutant and wild-type Calpain 3 complexes may be affected without inhibiting the wild-type activity per se. However, a more thorough investigation is necessary to understand the molecular mechanism and dominant inheritance of the heterozygous 21-bp deletion mutation in LGMDD4.

## INTRODUCTION

Limb-girdle muscular dystrophy (LGMD) is a group of neuromuscular diseases characterized by progressive muscle weakness and muscular atrophy of the proximal muscles caused by a defect in Calpain 3^1^. Calpain 3 is a member of the Calpain protease family, which are non-lysosomal calcium-regulated proteolytic enzymes involved in various biological phenomena. Among all calpain isoforms reported in humans, the Calpain 3 (CAPN3) isoform is predominantly found in the skeletal muscles^2^. Calpain 3 forms an integral part of the sarcoplasmic reticulum of the skeletal muscles and plays a role in maintaining muscle integrity and function by regulating sarcomeric protein turnover and maintaining the integrity of the sarcomere structure^3,4^.

The most common form amongst all Calpainopathies is Limb Girdle Muscular Dystrophy Recessive 1(LGMDR1), formerly known as Limb Girdle Muscular Dystrophy type 2A(LGMD2A), caused by a recessive mutation in the Calpain 3 (CAPN3) gene. The major clinical symptoms were characterized by symmetric and progressive weakness of proximal and pelvic limb girdle muscles and joint contractures with elevated creatine kinase levels in most of the cases ^5^. LGMD is primarily thought to be inherited as a recessive trait, but little evidence suggests its transmittance through an autosomal dominant inheritance pattern in several generations^6,7^. For the first time in 2016, Vissing and colleagues reported 37 patients from 10 European families with an autosomal dominant LGMD co-segregating with an in-frame 21-bp deletion (c.643_663del21, p. Ser215_Gly221del) in CAPN3^8^. Nine affected patients had a milder phenotype than those affected by LGMDR1, with normal expression of the mutated mRNA and no evidence of nonsense-mediated mRNA decay, but there was a significant decrease (<15% of control values) at protein levels measured by Western blot analysis. Therefore, the authors proposed a dominant-negative effect of the CAPN3 deletion through inhibition of wild-type Calpain 3 activity by mutant Calpain 3 when they form homomeric complexes^9^. Another group also reported the autosomal dominant inheritance of LGMD with the CAPN3 21-bp deletion independently ^9^. In that study, three unrelated individuals of North European origin with heterozygous 21-bp deletion mutation showed reduced muscular Calpain 3 protein levels ^9^. In this study, we sought to understand the phenomenon of the dominant toxicity of the 21-bp deletion mutation of Calpain 3 by examining catalytic activity and dominant phenotypic aberration using a human cell lines and *Drosophila*, respectively.

## METHODS

### DNA constructs

The human Calpain 3 and 21 bp-deletion Calpain 3 mutant were synthesized and sub-cloned in the pcDNA3.1 vector containing a C-terminal FLAG tag by GenScript. Wild-type and deletion mutant plasmid constructs for *Drosophila* Calpain A and Calpain B were synthesized and sub-cloned in the pUASTattb vector to generate transgenic *Drosophila* lines from BestGene.

### Cell culture and transfection

HeLa cell lines were purchased from the American Type Culture Collection (ATCC). Cells were maintained in Dulbecco-modified Eagle medium (Gibco) supplemented with 10% fetal bovine serum (Gibco), 1× penicillin/streptomycin (Invitrogen), and 1 x GlutaMax (Gibco). Cells were transfected using jetPRIME (Polyplus) transfection reagent according to the manufacturer’s protocol.

### Immunoblotting and Antibodies

Cell samples were collected and lysed in ice-cold RIPA buffer (Cell Signaling Technology) with protein inhibitor cocktail (Sigma) and subsequently separated by sodium dodecyl sulfate-polyacrylamide gel electrophoresis (SDS-PAGE) after measuring protein concentration by bicinchoninic acid (BCA) (Pierce). The following primary antibodies were used: FLAG (Sigma and Proteintech, 1:1000), Calpain 3 (Proteintech and Santa Cruz Biotechnology, 1:1000), Calpastatin (Proteintech and Santa Cruz Biotechnology, 1:1000), Actin (Proteintech and Santa Cruz Biotechnology, 1:5000). Immunoblots were visualized and analyzed with the Odyssey FC System (LI-COR).

### Generation of Drosophila lines

Transgenic *Drosophila* lines were generated by BestGene with the standard PhiC31 integrase-mediated transgenesis system. All transgenes were inserted into the third chromosome attp2 site to avoid positional gene expression differences. Fly cultures and crosses were performed on standard fly food (Genesee Scientific) and raised at 25°C with a 12:12 hr light:dark cycle. GMR-GAL 4 and MHC-GAL4 were used as drivers for their expression in the eye and muscles, respectively.

### Drosophila adult muscle preparation and immunohistochemistry

The muscle integrity and sarcomere structure of the indirect flight muscle were analyzed as previously described^10^. Briefly, adult flies (4-week-old) were quickly dissected and fixed with 4% paraformaldehyde in 1X PBS for 1 hour, embedded in OCT compound (Fisher Scientific), and snap frozen in liquid nitrogen or dry ice. 15-16 microns thick sections were cut by a cryomicrotome (Leica) and directly mounted on the slide. Additional fixing was performed with 4% paraformaldehyde in 1XPBS for 10 minutes at room temperature prior to permeabilization with 0.2% Triton X-100 buffer in PBS and blocked with 5% BSA solution in PBS for 1 hour. Sections were incubated with phalloidin–Alexa Fluor 594 (Invitrogen) overnight at 4°C for muscle staining and finally were mounted with Prolong Diamond Antifade Reagent with DAPI and imaged with an LSM 710 confocal microscope (Carl Zeiss) with 63× magnification.

### Statistical Analysis

Graphs for the quantitative analysis of immunoblots (mean difference between groups with 95% confidence interval) were generated using Estimation Stats, http://estimationstats.com^11^. We performed an unpaired T-test for the evaluation of our data with a two-sided permutation p-value.

## RESULTS

### The deletion mutant of Calpain 3 loses its autocatalytic activity

To test the expression and stability of wild-type and deletion mutant Calpain 3, we transiently transfected C-terminal FLAG-tagged wild-type (WT) and 21-bp deletion mutant (MT) Calpain 3 plasmid constructs to HeLa cells (which is not expected to express Calpain 3) followed by immunoblotting (Fig. 1A). Unexpectedly, the overexpression of FLAG-tagged wild-type (WT) was not detected in the immunoblot by anti-FLAG antibody, but there was a prominent expression band of FLAG-tagged deletion mutant protein (MT) (Fig. 1B, left panel). In contrast, a faint band close to the molecular weight of Calpain 3 (94 kDa) was detected in the Empty Vector (EV) control lane and wild-type (WT) when immunoblotted with anti-Calpain 3 antibody. However, we speculate that these bands were non-specifically detected by anti-Calpain 3 antibody according to a negligible amount of mRNA level in HeLa cells reported in the human protein atlas (https://www.proteinatlas.org/). Consistent with the immunoblotting result with anti-FLAG antibody, the anti-Calpain 3 antibody detected an intense band of transiently transfected deletion mutant Calpain 3 (MT) (Fig. 1B, right panel). This result can be deciphered by the unique characteristic of Calpain 3 proteins. Calpain 3 undergoes extremely rapid and exhaustive autolysis dependent mostly upon IS1 and IS2 regions within Calpain 3 (Fig. 1 A)^4,12^. Thus, transiently transfected Calpain 3 proteins were rapidly auto-degraded and could not be detected by anti-FLAG and anti-Calpain 3 antibodies (Fig. 1 A and B). However, the loss of autolytic activity in the deletion mutant form (MT) produced stable proteins detected by anti-FLAG and anti-Calpain 3 antibodies (Fig. 2B, right panel). To confirm the autolytic activity of wild-type and mutant Calpain 3, we examined the intermediate autolyzed fragments in this system. Typically, two fragments are generated upon degradation around 30 kDa and 60 kDa. Since we used C-terminal FLAG-tagging, a smaller N-terminal autolyzed fragment (∼30 kDa) was detected by anti-Calpain 3 antibody (Fig. 1A), and a larger C-terminal autolyzed fragment (∼60 kDa) was detected by anti-FLAG antibody in the WT lane (Fig. 1C and Suppl. Fig. 1). A full-length deletion mutant Calpain 3 (MT) was very well expressed when probed with both anti-FLAG and anti-Calpain 3 antibodies but no autolyzed fragments were detected (Fig. 1C). Thus, the 21-bp deletion mutant protein is an inactive form of Calpain 3 for its autolysis.

**Figure 1.**
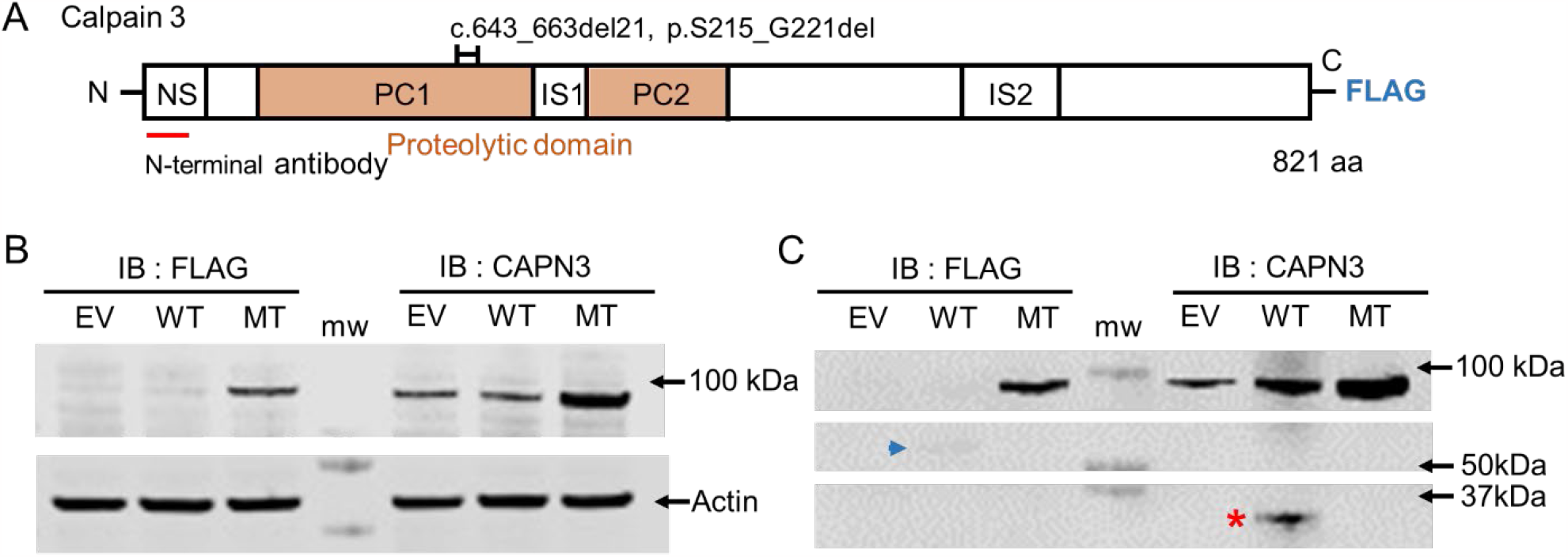
The deletion mutant Calpain 3 loses its autocatalytic activity. HeLa cells were transfected with FLAG-tagged wild-type Calpain 3 (WT), 21-bp deletion mutant (MT) Calpain 3, and empty vector (EV). (A) Schematic diagram of *C*-terminal FLAG-tagged Calpain 3 and deletion mutation, a 21-bp (7 amino acids) deleted region, and an *N*-terminal anti-Calpain 3 antibody binding site (marked in a black bracket and a red line), respectively. NS: N-terminal leader Sequence, PC1 & 2: Protease Core1 & 2, IS1 & 2: Insertion Sequence1 & 2, (B) Immunoblotting with anti-FLAG, anti-Calpain 3, and anti-Actin (loading control) antibodies on overexpression of Empty vector (EV), wild-type (WT), and mutant (MT) Calpain 3 in HeLa cells. FLAG-tagged MT was detected but not WT (Left panel). An increased MT band was detected by but not WT (Right panel), (C) Immunoblotting for detection of autolyzed fragments of Calpain 3. FLAG-tagged Wild-type Calpain 3 (WT) was rapidly processed by its own autocatalytic activity (blue arrowhead indicates the C-terminal larger fragment and red asterisk indicates the smaller N-terminal fragment) but none in the case of the mutant lane (MT).

**Figure 2.**
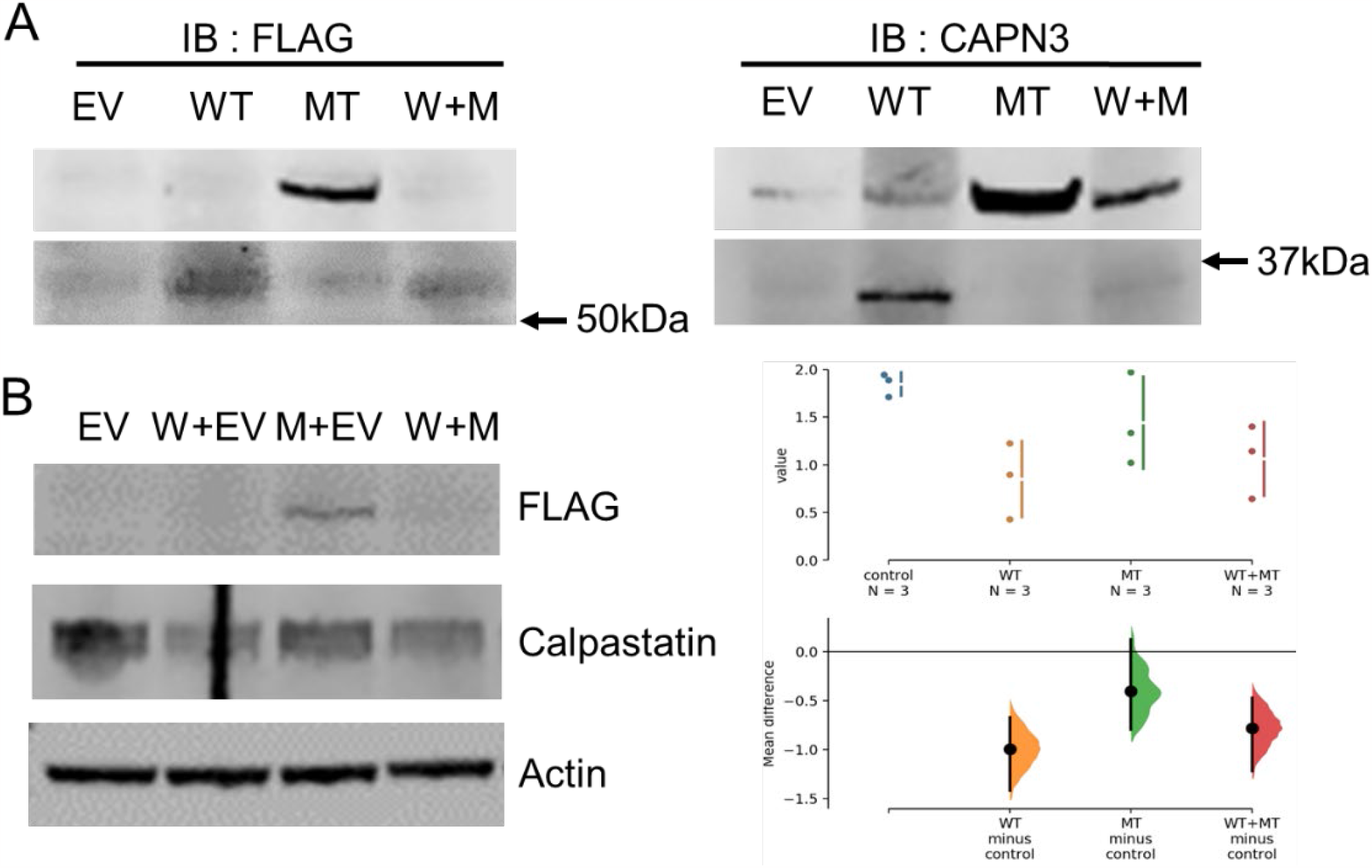
The deletion mutant Calpain 3 does not inhibit wild-type activity. Hela cells were co-transfected with wild-type (WT) and mutant (MT). Immunoblotting was performed by anti-FLAG and anti-Calpain 3 antibodies. (A) Left panel: Wild-type Calpain 3 was normally autolyzed in both lanes in (WT) alone and also along with the deletion mutant co-expression lane (W+M). Right panel: The mutant proteins were also normally processed and detected by anti-Calpain 3 antibody. (B) Immunoblot for Calpastatin expression detected by anti-Calpastatin antibody. Calpastatin was also normally processed with wild-type/mutant co-expression at a comparable level to wild-type overexpression. (Left panel. Quantification with three independent replicates (Right Panel, two-sided permutation p values: 0.0, 0.29, 0.0 respectively).

### The deletion mutant Calpain 3 does not inhibit wild-type activity

Calpain 3 is believed to form a homodimer or a homotrimer ^13,14^. Since mutant Calpain 3 is auto-catalytically inactive, mutant proteins may dominant-negatively interfere with the activity of this homomeric complex, as proposed by Vissing et al. ^8^. Therefore, we tested this hypothesis by co-transfecting with both wild-type and deletion mutant (MT) constructs. We examined whether mutant Calpain 3 inhibited the autolytic activity of the wild-type Calpain 3 and also its catalytic activity against Calpastatin, which has been known to be degraded by Calpain 3 ^15,16^. As seen in Fig. 2A, normal autolysis of wild-type Calpain 3 was observed in the wild-type and deletion mutant co-expression panel (W+M). Corresponding fragments were also detected with anti-FLAG and Calpain 3 antibodies (Fig. 2A, lower, Suppl. Fig. 2), as expected. Interestingly, when the mutant protein was co-expressed with wild-type protein, the expression of Calpain3 was decreased significantly compared to when mutant protein was expressed alone (Fig. 2A). This indicates that inactive mutant Calpain 3 was normally degraded by wild-type Calpain 3 and the autocatalytic activity of wild-type Calpain 3 was not significantly reduced by the inactive Calpain 3 mutant. When we examined the levels of Calpastatin as a readout of wild-type Calpain 3 activity to reveal the relationship between wild-type and mutant Calpain 3, Calpastatin was decreased by the overexpressed wild-type Calpain 3 (W+EV) compared to the empty vector-transfected control (Fig. 2B). When mutant Calpain 3 was overexpressed, there was no change in the Calpastatin level. When wild-type and mutant were co-expressed (W+M), the Calpastatin level was similarly decreased to that observed in the wild-type (W+EV) lane, suggesting that the mutant form could not block the wild-type catalytic activity significantly (Fig. 2B). This phenomenon was observed in three independent replicates and was statistically significant (Fig. 2B, right panel, Suppl. fig 3). These results, altogether, demonstrated that the deletion mutant did not inhibit wild-type Calpain 3 autolysis and catalytic activity against another substrate. In addition, the deletion mutant protein can be normally processed by wild-type Calpain 3 intermolecularly.

### The deletion mutants of *Drosophila* Calpain A and B do not have dominant toxicity in the *Drosophila* eye

We then explored the effect of wild-type and mutant Calpain 3 using an *in vivo* fly model. We generated a *Drosophila* version of the 21-bp deletion on the two *Drosophila* homologs of human Calpain 3, Calpain A and Calpain B, and their deletion mutant counterparts, delCalpain A and delCalpain B^17,18^. To test whether the overexpression of these constructs could cause dominant toxicity, we first expressed Calpain A, delCalpain A, Calpain B, and delCalpain B in *Drosophila* eyes, which are not expected to express endogenous Calpain A and B, with the glass multimer reporter (GMR)-GAL4 driver. All of them did not cause any significant abnormal eye phenotypes at eclosion and at four weeks old (Fig 3). These findings indicate that the dominant toxicity of the 21-bp deletion cannot be recapitulated in *Drosophila* eyes and also suggest that the 21-bp deletion mutant may not act in a dominant gain-of-toxicity mechanism.

**Figure 3.**
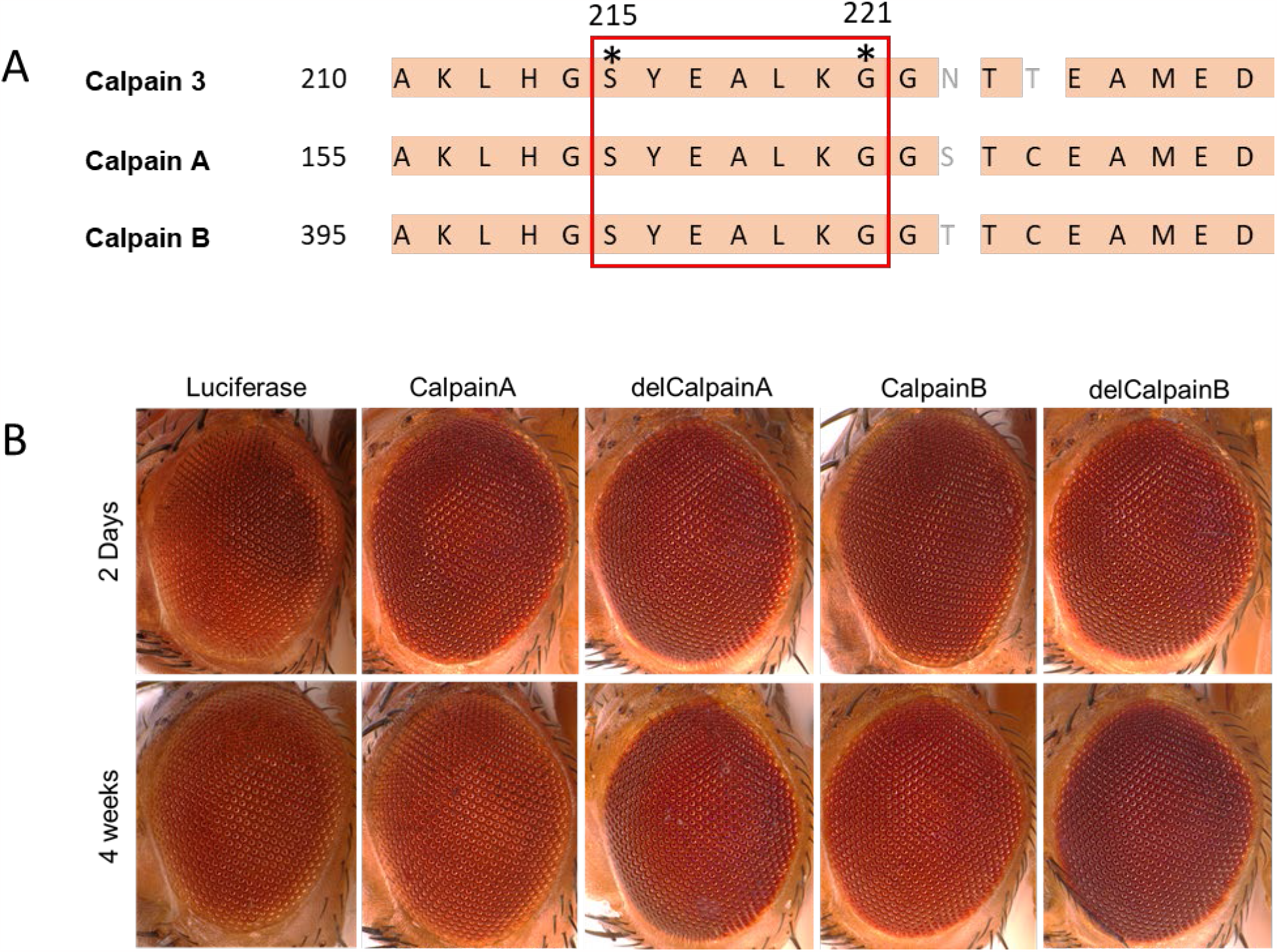
The deletion mutants of *Drosophila* Calpain A and B do not have dominant toxicity in the *Drosophila* eye. (A) Amino acid sequence alignment around 21-bp deletion region across Human Calpain 3 and *Drosophila* Calpain A and B. The conserved sequences are shown relative to the positions of the translation start codon (indicated by bold letters with colored backgrounds). All the seven amino acids in the deleted regions are highly conserved across the species (marked with a red box). (B) Wild-type and mutant Calpain A and B were expressed in *Drosophila* eyes with GMR-GAL4. In 2 days (upper panel) and four weeks-old flies (lower panel), no significant developmental and age-related dominant toxicity were observed (females, at 25 °C).

### The deletion mutants of *Drosophila* Calpain A and B did not show dominant toxicity in indirect flight muscles

Calpain 3 is an integral part of skeletal muscle cells, and muscle-specific activities of Calpain A and B have been reported in *Drosophila*.^19^ Therefore, to better investigate the function of mutant Calpain 3 that may act in a dominant-negative manner, we expressed Calpain A, delCalpain A, Calpain B, and delCalpain B in *Drosophila* muscles using the Myosin Heavy Chain (MHC)-GAL4 driver. Immunohistochemical analysis of thoracic indirect flight muscle sections of flies overexpressing wild-type and mutant Calpain A and B did not show significant muscle degeneration or sarcomere structure abnormality in 4-week-old aged flies (Fig 4). Therefore, we did not observe any dominant toxicity of mutant Calpain 3, whether dominant gain-of-toxicity or dominant-negative, in our model systems.

**Figure 4.**
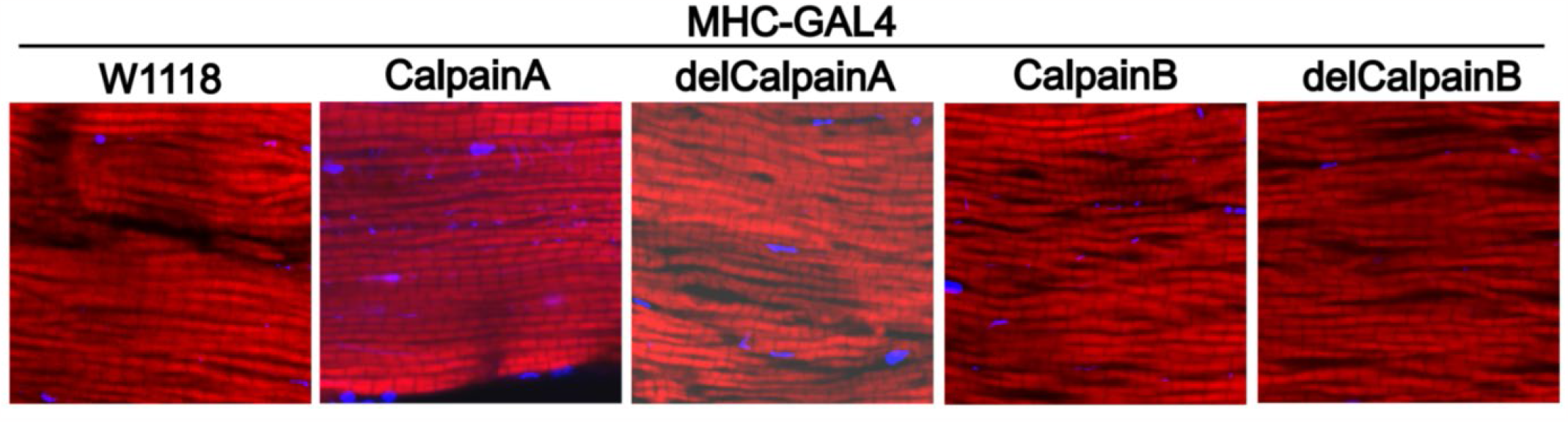
The deletion mutants of Drosophila Calpain A and B did not show dominant toxicity in indirect flight muscles. Wild-type and mutant Calpain A and B were expressed in *Drosophila* muscles with MHC-GAL4. Representative immunohistochemical images of the thoracic segment of four-week-old flies showing no significant muscle degeneration and related abnormalities. (Red: Phalloidin, myofibrils, Blue: DAPI, nucleus).

## DISCUSSIONS

We have provided here experimental evidence to show that the 21-bp in-frame deletion mutation of Calpain 3 causes loss of autolysis activity, and the mutant form does not inhibit the normal catalytic activity of its wild-type counterpart. In addition, the mutant protein can be normally processed by the wild-type Calpain 3. Although we tried to detect the dominant toxicity of the mutant protein by overexpression of the mutant Calpains in the *Drosophila* eye and muscle, we did not observe any significant dominant toxicity in both tissues.

Vissing and colleagues proposed the dominant-negative mechanism for the LGMDD4 with a 21-bp deletion mutant according to its inheritance pattern, molecular evidence showing normal Calpain 3 mRNA level without nonsense-mediated decay, and compromised protein levels of Calpain 3 in the muscle biopsies. Although other potential possibilities explaining the dominant inheritance of the 21-bp deletion mutation in LGMDD4 have been suggested, such as a possibility of a second mutation associated with CAPN3 gene or acting as an exonic enhancer, and co-transmitted secondary mutation in a different gene ^20^, Vissing et al. reasonably addressed those concerns ^21^ and an additional report from an independent group further reduced the concerns ^9^. Vissing et al. proposed a dominant negative mechanism via the inhibition of Calpain 3 protein activity by the deletion mutant Calpain 3 ^8^. However, our immunoblot data from HeLa cells suggest that the dominant negative effect may not be dependent on inhibiting the wild-type Calpain 3 activity per se by the mutant form. Our results strongly support that the impaired stability of Calpain 3 homomeric complexes may be the key to understanding the decreased Calpain 3 levels (< 15%) in patients. Calpain 3 protein was reported to form a trimeric protein recently^14^. If only wild-type homotrimeric complexes are stable and others containing a mutant subunit are not, then only 12.5% of Calpain 3 can be stably maintained in patients. This corresponds well with the levels of Calpain 3 in patients. A detailed biochemical analysis is required to understand how mutant Calpain 3 affects the stability of complexes without affecting the wild-type activity itself.

Unfortunately, however, we did not observe any dominant toxicity in our *Drosophila* models using two different Calpain 3 mutants, whether dominant gain-of-toxicity or dominant negative. This may imply that the impact of the deletion mutant Calpain 3 should be examined at other developmental stages of muscle differentiation along with other drivers, since MHC-GAL4 drives transgene expression in differentiated muscles. In addition, it may be necessary to study the effect of the mutant Calpain 3 using human myoblasts or differentiated myotubes to gain more insight. Nevertheless, our data reveal the important properties of the mutant Calpain 3 produced by the 21-bp deleted gene and exclude a possible dominant-negative mechanism by inhibiting the wild-type activity per se, and suggest a possibility of dominant negative effects via the decreased stability of complexes or the active degradation of complexes through an unknown quality control mechanism. It may be possible that the protein complex formed with the wild-type and mutant Calpain 3 is less stable and degraded quickly in patients. However, a more precise investigation is necessary to elucidate how the patients have reduced Calpain 3 protein level with a normal mRNA level in LGMDD4.

## Abbreviations

AD: autosomal dominant
CAPN3: calpain-3
CK: creatine kinase
LGMD: limb-girdle muscular dystrophy
LGMDD4: autosomal dominant limb-girdle muscular dystrophy type
LGMDR1: recessive limb-girdle muscular dystrophy type
WT: wild-type
MT: mutant
EV: Empty Vector.

## Acknowledgment

We want to thank Maddie Chalmers for her important feedback during manuscript preparation. We are also thankful to Minwoo Baek for helpful technical guidance during the experiments. We appreciate Drs. Zhiyv Niu and Margherita Milone for their insightful discussions.

## Author contributions

N.C.K. conceived the project. Y.-J.C, S.J.O, S.M and J.H.K designed and performed the experiments. Y.-J.C, S.J.O, S.M analyzed data and S.M, S.J.O, and N.C.K wrote the manuscript.

## Conflicts of Interest

The authors have no conflicts of interest to disclose.

**Supplemental Figure 1.**
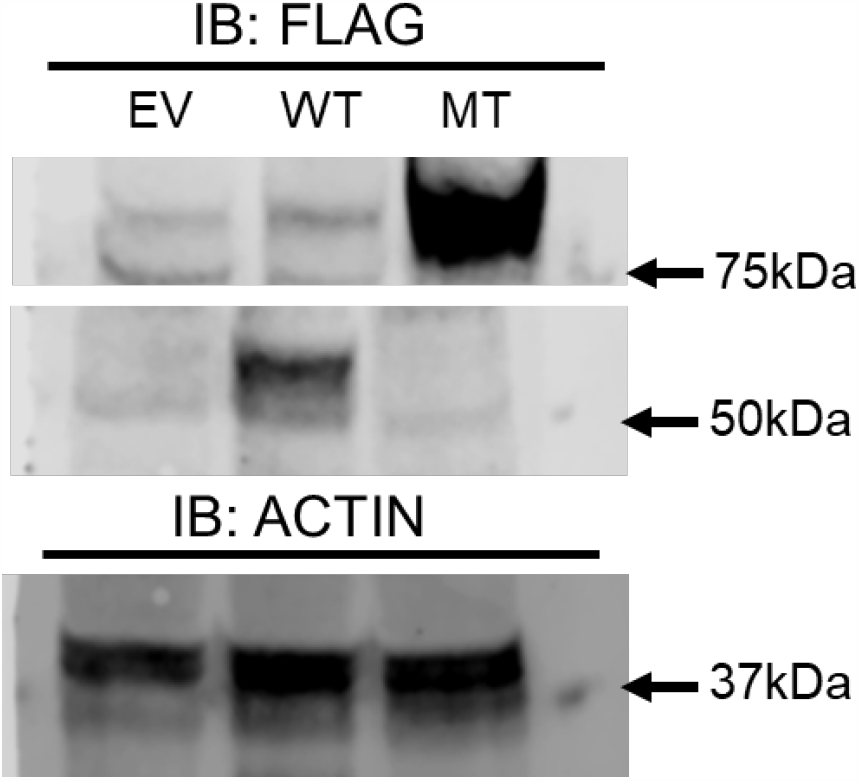
Immunoblotting for detection of autolyzed fragments of Calpain 3. FLAG-tagged C-terminal fragment of Wild-type Calpain 3 was clearly visible when probed with anti-FLAG antibody but none in the case of mutant lane (MT). Actin was kept as a loading control.

**Supplemental Figure 2.**
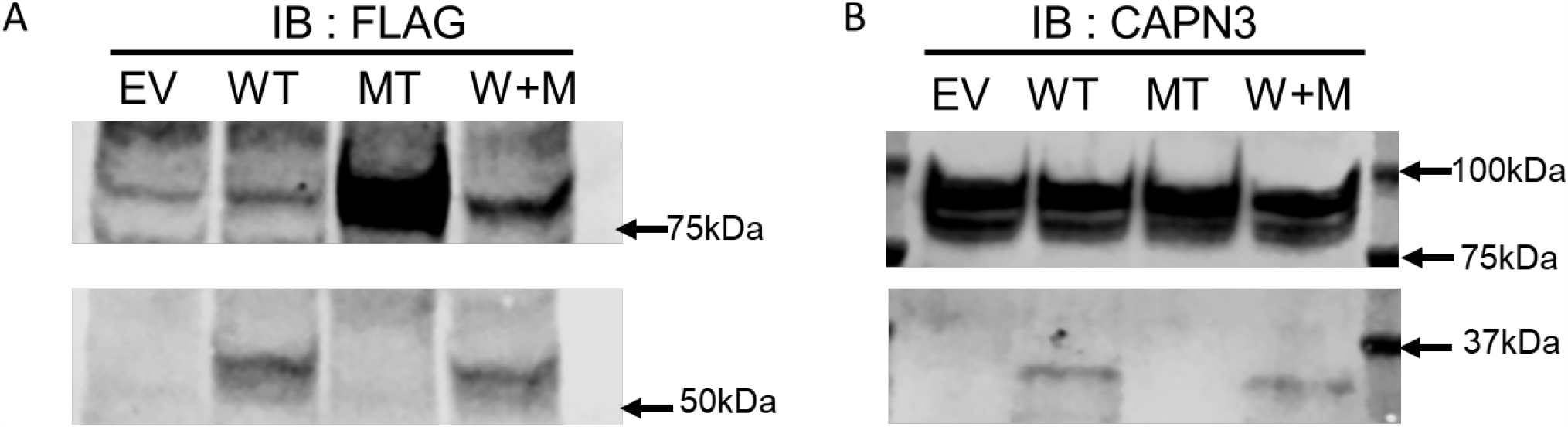
Immunoblots showing the detection of N-terminal (A) and C-terminal (B) autolyzed fragments of Calpain 3 in Empty vector (EV), Calpain 3 wild-type (WT), and Calpain 3 deletion mutant (MT) transfected HeLa cell lysates. Please note that this (B) was an independent experiment from (A) and the blot was overexposed in order to have a clear view of the hard-to-detect smaller fragments in WT and W+M lanes.

**Supplemental Figure 3.**
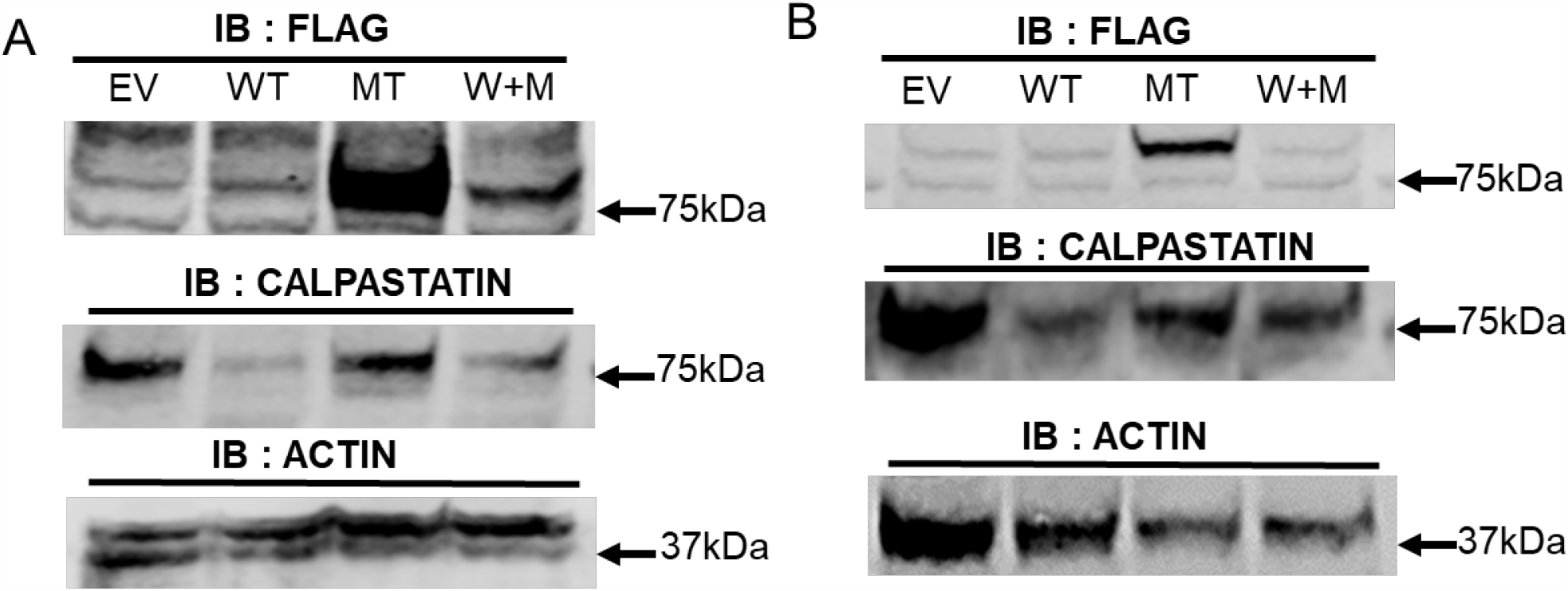
Immunoblotting for detection of Calpastatin expression in Empty vector (EV), Calpain 3 wild-type (WT), and Calpain 3 deletion mutant (MT) transfected HeLa cell lysates. Actin was kept as a loading control (Quantification and statistical results are in Figure 3B).

